# Neuronal receptors display cytoskeleton-independent directed motion on the plasma membrane

**DOI:** 10.1101/269381

**Authors:** R. D. Taylor, M. Heine, N. J. Emptage, L. C. Andreae

## Abstract

Directed transport of transmembrane proteins is generally believed to occur via intracellular transport vesicles. However, using single particle tracking in rat hippocampal neurons with a pH-sensitive quantum dot probe which specifically reports surface movement of receptors, we have identified a subpopulation of neuronal EphB2 receptors that exhibit directed motion between synapses within the plasma membrane itself. This receptor movement occurs independently of the cytoskeleton but is dependent on cholesterol and is regulated by neuronal activity.

## Introduction

Membrane proteins are known to exhibit lateral diffusion within the plasma membrane (Borgdorff & Choquet, 2002; Dahan et al., 2003; Heine et al., 2008), while their directed transport is believed to occur via the intracellular movement of transport vesicles. These endosomal transport vesicles travel along the cytoskeletal network of the cell, driven by molecular motors (Hirokawa & Takemura, 2005; Kapitein et al., 2010). This transport is especially important in polarized cells such as neurons, where, for example, the rapid delivery of key receptors into and out of synapses can be critical for neuronal function. Although directed movement of GABA_A_ receptors has been described in neuronal growth cones (Bouzigues, Morel, Triller, & Dahan, 2007) this was completely dependent on the microtubule network, and given that quantum dot based probes are able to report individual receptors switching between surface diffusion and intracellular active motor transport (Vermehren-Schmaedick et al., 2014), a confident description of movement on the membrane surface requires surface-specific protein tracking. Here, we use such an approach to track the transmembrane tyrosine kinase receptor, EphB2, a well-known neuronal protein with important roles in synapse formation (Sheffler-Collins & Dalva, 2012) and plasticity (Contractor et al., 2002; Grunwald et al., 2001).

## Results

In order to specifically target single molecule imaging to the cell surface, we developed a probe to differentiate between receptor movement on the neuronal cell surface versus that within endosomal transport vesicles. The intraluminal (internal) environment of intracellular vesicles is relatively acidic (pH<6) (Ouyang et al., 2013). We exploited our observation that the level of fluorescence emitted by the 655nm wavelength quantum dot (QD) exhibited significant pH sensitivity. We labelled endogenous EphB2 receptors in cultured neurons with the QD-655 conjugate and examined the effect of changing the external pH. Bath application of pH6 solution resulted in a dramatic quenching of QD fluorescence intensity (Figure 1A) to a level where QDs could no longer be detected (Figure 1B), while pH7 solution had no effect. Since QD655 fluorescence is virtually undetectable at pH6, this suggested that any visible QDs should reside on the neuronal surface as those lying within vesicles would be effectively invisible. To verify that this was indeed the case, we used the membrane impermeable dye, QSY-21, which is known to quench QD fluorescence (Jablonski, Kawakami, Ting, & Payne, 2010). Addition of QSY-21 resulted in the inability to detect any QD-EphB2Rs within 10s of its application (Figure 1C); further indicating that all visible QD-EphB2Rs are on the cell surface.

**Figure 1.**
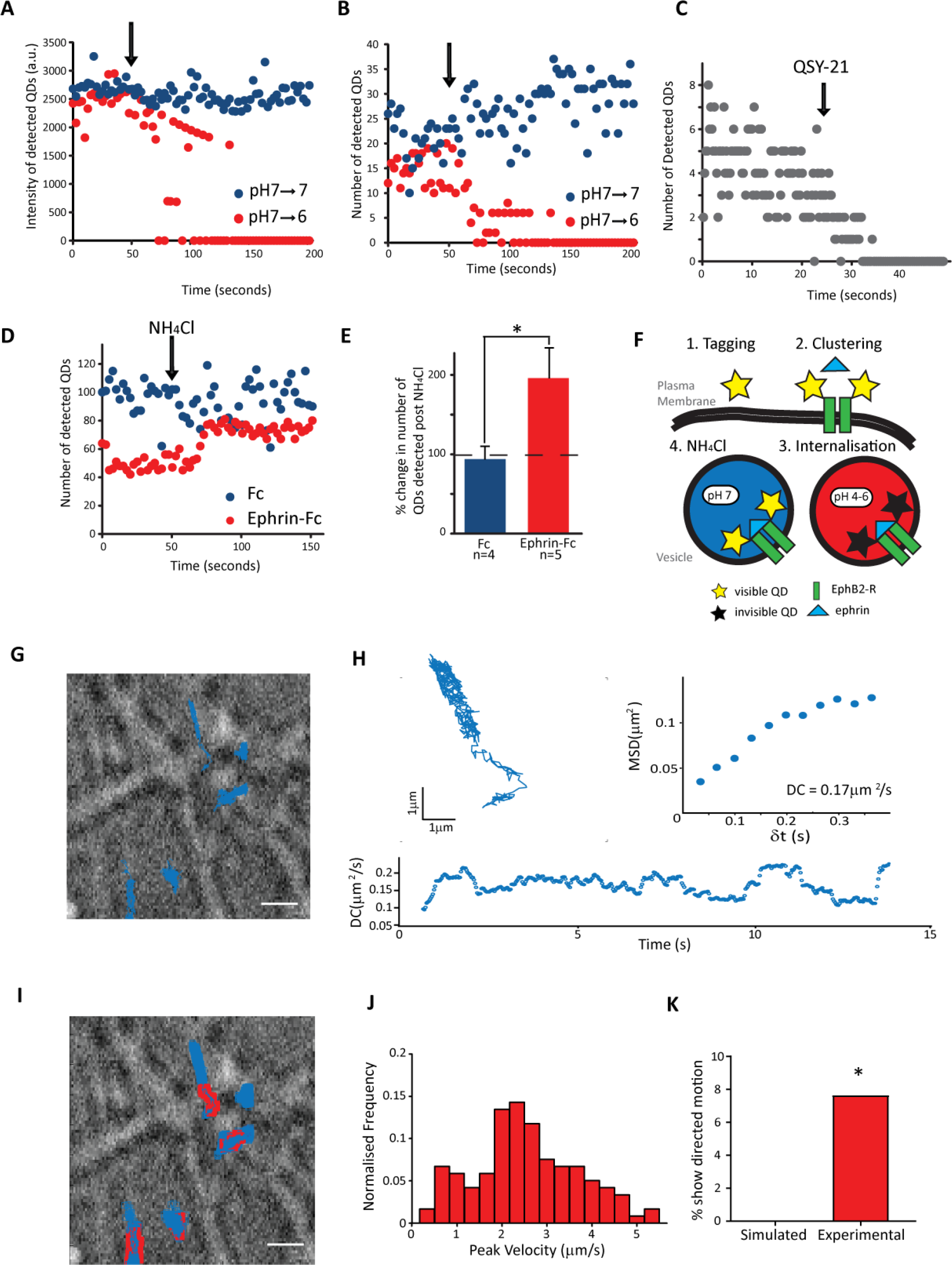
Single particle tracking of EphB2Rs in hippocampal neurons with a surface specific quantum dot probe reveals directed motion. (**A-D)** Arrows indicate addition of novel solutions. Mean fluorescence intensities (**A**) and total number of detected single molecules (**B**) of QD-EphB2Rs following addition of pH6 solution (red circles) compared with pH7 control (blue circles). (**C**) Number of detected QD-EphB2Rs following addition of QSY-21 (100μM) (**D-E**) Dissipation of intracellular pH gradients by application of NH_4_Cl (50mM, pH7.2) to neurons previously incubated with pre-clustered ephrinB1 (Ephrin-Fc; red circles, n=5) or control Fc fragments (Fc; blue circles, n= 4) leads to a significant increase in detected QD-EphB2Rs, (**D**) population averages, (**E**) quantification (p<0.05, graph shows mean±S.E.M). (**F**) Model illustrating experiments shown in (**A**) and (**C**): Clustering of QD-labelled EphB2Rs leads to receptor internalisation and hence QD fluorescence quenching. These QD-EphB2Rs are then revealed by neutralization of pH. (**G**) Example trajectories of QD-EphB2Rs (blue) along neuronal processes. Scale bar represents 10μm. (**H**) MSD plotted against time interval (δt) for illustrated trajectory yields a diffusion coefficient (DC) of 0.17μm^2^/s by linear fit, while DC is seen to vary over the trajectory by time resolved MSD analysis. (**I**) Trajectories from (**G**) showing episodes of directed motion in red. (**J**) Frequency distribution of detected peak velocities in experimental data (n=119; mean = 2.51±1.16μm/s. (**K**) Quantification to compare percentage of experimental QDs showing directed motion with simulated data of Brownian motion, (Chi-squared, p<0.0001). See also Figure S1.

To confirm that intravesicular QD-labelled EphB2Rs are not detectable we pre-treated neurons with clustered ephrinB1-Fc, a high-affinity ligand for EphB2Rs which has been shown to cluster EphB2Rs and cause internalization to endosomes (Zimmer, Palmer, Kohler, & Klein, 2003). If internal QD-EphB2Rs are present, but rendered undetectable by the relatively acidic endosomal environment, we reasoned that they should be revealed by neutralizing the endosomal pH gradient with ammonium chloride (NH_4_Cl) (Axelsson et al., 2001). Upon addition of NH_4_Cl we saw a significant increase in the number of detected QD-EphB2Rs, compared with neurons that were treated with control Fc only (Figure 1D-F). This unveiling of internalised QD-EphB2Rs further confirms that QD-EphB2Rs travelling within vesicles inside the cell are not visible. Taken together, these experiments demonstrate that all visible QD-EphB2Rs are located on the cell membrane surface.

Single particle tracking (SPT) of QD-EphB2R trajectories in hippocampal neurons revealed that single receptors are not static but move laterally in the neuronal cell membrane (Figure 1G and Supplemental Movie). Standard approaches to analysis of single particle movement have utilised the mean squared displacement (MSD) method to distinguish between three types of motion displayed by particles: confined diffusion, free Brownian diffusion and super-diffusive or ‘directed’ motion. This type of analysis generally assigns a single motion mode to each particle trajectory. However, we noticed that while some QD-EphB2Rs were either stationary or mobile for the duration of the recording, others interchanged between periods of relative localised stability and unrestricted motion (Supplemental Movie), suggesting that a more time-resolved analysis would be able to better describe individual trajectories. Indeed, segmenting trajectories by synaptic location has previously demonstrated that the diffusion characteristics of neurotransmitter receptors vary depending on whether they are located at or outside the synapse (Renner, Schweizer, Bannai, Triller, & Levi, 2012). We therefore analysed the trajectories of QD-EphB2Rs by calculating MSD within sliding time windows, based on an approach previously validated in microtubule transport (Arcizet, Meier, Sackmann, Radler, & Heinrich, 2008). The method determines two coefficients that describe the change in MSD with time interval within each window. The diffusion coefficient (DC) is calculated over the linear portion (i.e. the first few points) of the function. The exponent α describes the curvature of the function and thus is an indicator of the type of motion displayed by the receptor, where α=1 represents diffusion, <1 indicates confined diffusion and >1 directed motion, and it is resolved over a longer timescale. This approach confirmed our impression that individual receptors exhibit varying DCs over the time-course of a single trajectory (Figure 1H). Further, to our surprise, we found that some receptors appeared to undergo short periods of directed motion (71/1287) (Figure 1I) with an average peak velocity of 2.51±1.156 μm/s (Figure 1J). We compared experimental data with time-resolved analysis of simulated trajectories of randomly moving particles displaying Brownian motion. We found that there was a significantly greater proportion of experimental EphB2R trajectories exhibiting periods of directed motion than of simulated ones, indeed with no such episodes arising from analysis of the simulated pure diffusional trajectories (Experimental trajectories: 14/183 [8.44,22.75]; Simulated Brownian trajectories: 0/350 [0,3.8] Chi squared test, p<0.0001, Figure 1K). To determine whether it might be possible to detect apparent directed motion as a result of the geometry of membrane space available for diffusion, we modelled diffusional movement of particles in a variety of spatially defined regions. In no case were we able to identify directed movement (see Figure S1A,B). Reduced sampling rates also only reduced the apparent α (Figure S1C). Since all labelled EphB2Rs in our experiments are restricted to the external neuronal surface, such that the episodes of directed motion cannot be due to intracellular vesicular transport, these results demonstrate that these receptors have the capacity to move with directed motion on the plasma membrane.

We next asked to what extent either the diffusive or the directed motion of surface EphB2Rs is dependent on the cell cytoskeleton. We conducted QD-EphB2R tracking experiments in the presence of cytoskeletal modifying drugs and compared their motion characteristics with vehicle control (DMSO). When we examined diffusive motion, we found that there were two populations of diffusing QD-EphB2Rs: super-restricted (mean α = 0.026); and diffusing (mean α=0.59) (Figure 2A). Recently, an extension of the fluid mosaic model describing cell membrane structure (Singer & Nicolson, 1972) has been proposed which describes different compartments within a phospholipid-cholesterol sheet, supported by a meshwork of microtubules and actin filaments (Kusumi, Suzuki, Kasai, Ritchie, & Fujiwara, 2011). This model envisages membrane structure as being composed of an actin-based ‘fence’ and transmembrane protein ‘pickets’. We speculated that movement of the super-restricted group of receptors might be limited by this actin fence, and indeed, disruption of the actin cytoskeleton by addition of Latrunculin A (5μM) caused a significant increase in α (i.e. greater movement) of the super restricted group (Figure 2B) without affecting the more freely diffusing receptors (Figure 2C). No changes were seen following disruption of the microtubule network (Figure 2B,C). This is consistent with previous studies (Umemura et al., 2008) including of other neuronal receptors (Rust et al., 2010). However, neither modulation of the actin cytoskeleton (depolymerisation with Latrunculin A or stabilisation with Jasplakinolide) nor the microtubule network (depolymerisation with high dose Nocodazole or inhibition of microtubule dynamics with Nocodazole at low dose (Jaworski et al., 2009)) altered the directed motion characteristics of QD-EphB2Rs (Figure 2D-E).

**Figure 2:**
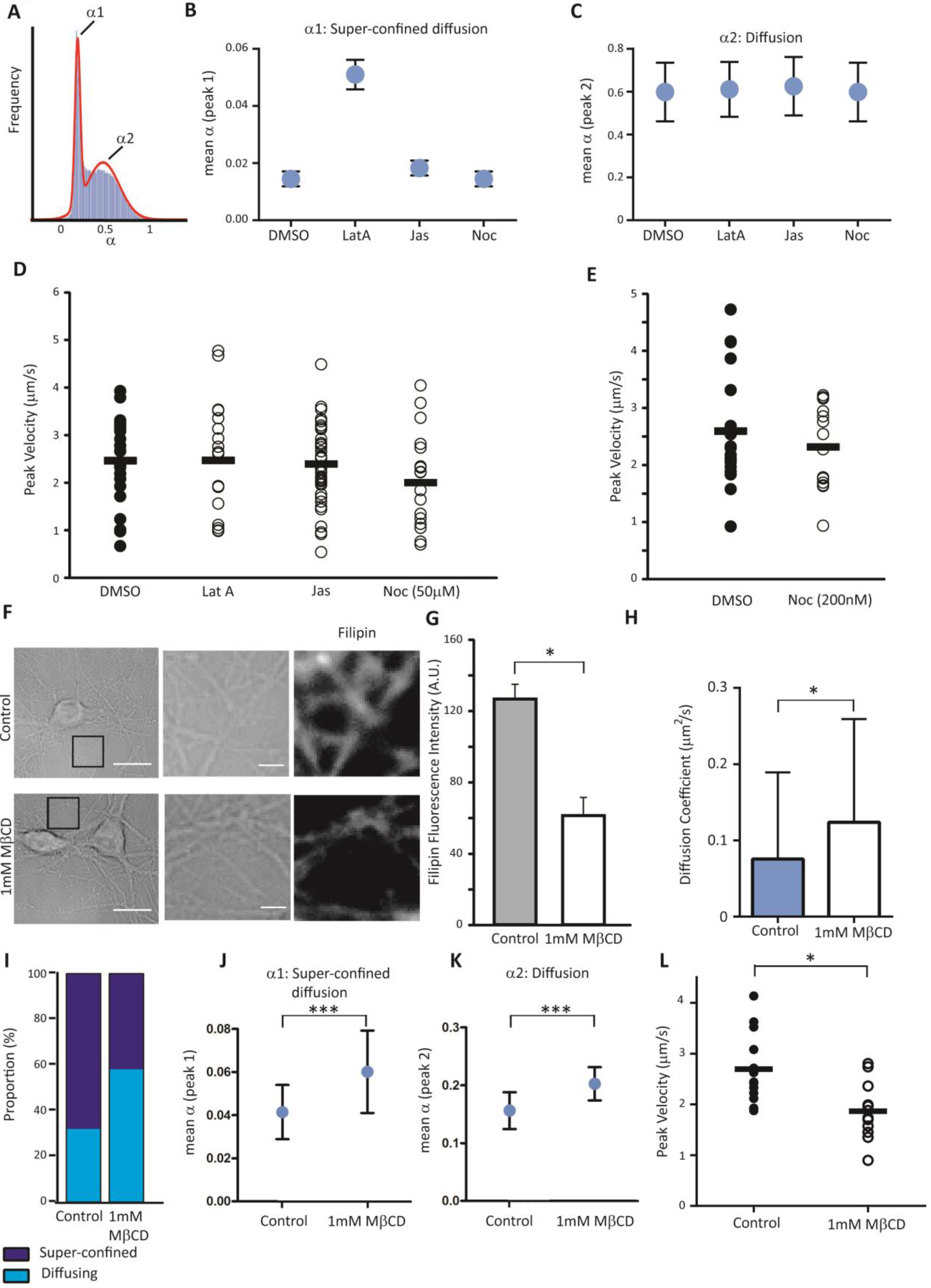
Directed motion of surface EphB2Rs is not affected by cytoskeletal modifying drugs but is dependent on membrane cholesterol. (**A**) Bimodal frequency distribution plot for values of instantaneous α in diffusing QD-EphB2Rs indicates a super-confined population (α_1_) and a more freely diffusing population (α_2_). (**B, C**) The effect of modulating the actin cytoskeleton (Lat A: Latrunculin A, 5μM; Jas: Jasplakinolide, 5μM) or the microtubule network (Noc: Nocodazole, 50μM) on α_1_(**B**) and α_2_(**C**). Error bars represent standard deviation. (**D**) Modifying the actin cytoskeleton or the microtubule network does not change the peak velocity of QD-EphB2Rs, bars indicate the mean, n =18-40. (**E**) as for (**D**), cells incubated in low dose Nocodazole. (**F-G**) Treatment with 1mM methyl-β-cyclodextrin (MβCD) reduces cholesterol levels; (**F**) left panels show representative brightfield images of dissociated neurons, scale bar 10μm, with higher power views of brightfield images (middle) and filipin fluorescence (right) of selected regions, scale bar 2μm; (**G**) Quantification of filipin fluorescence intensities, error bars represent SEM (control: n=48 dendrites, MβCD: n=49, p<0.05). (**H**) Cholesterol depletion with MβCD significantly increases the instantaneous DC of diffusing QD-EphB2Rs (control: 0.075μm^2^/s, M/CD:0.123 μm^2^/s, p<0.05, (**I**) MβCD treatment changes the relative proportions of diffusing QD-EphB2Rs that display super confined versus freely diffusing characteristics as defined by the value of instantaneous α, increases mean α_1_ (**J**) and to a lesser degree, α_2_ (**K**). (**H-K**) Error bars represent standard deviation. (**L**) Peak velocity of QD-EphB2Rs is significantly reduced by MβCD (n=13-17, p<0.05, Chi-squared).

We therefore hypothesised that the directed motion of EphB2Rs might be dependent in some way on the neuronal plasma membrane itself. Application of 1mM methyl-β-cyclodextrin (MβCD) resulted in depletion of membrane cholesterol in neuronal processes by approximately 50%, as quantified by levels of the macrolide Filipin, a cholesterol-binding fluorescent marker (Hering, Lin, & Sheng, 2003; Maxfield & Wustner, 2012; Renner, Choquet, & Triller, 2009) (Figure 2F,G). As previously reported (Renner et al., 2009), cholesterol depletion with MβCD resulted in an increase in DC that was accompanied by a change in the distribution of α (Figure 2H-K). While cholesterol depletion did not affect the proportion of QD-EphB2Rs displaying directed motion, it resulted in a significant reduction in the peak rate of directed motion. This indicates that the directed movement of surface EphB2Rs is dependent on membrane cholesterol.

We then questioned whether the directed motion of EphB2Rs exhibited any spatial specificity. A key feature of the neuronal landscape is the presence of postsynaptic specialisations at intervals along dendrites, which are known to impact receptor diffusion. For example, it has previously been shown that glutamatergic receptors show restricted diffusion within synapses and anomalous diffusion outside synapses (Groc et al., 2004; Heine et al., 2008). Using traditional immunostaining, we find EphB2Rs to be predominantly localized to dendrites, largely between synapses, in these cultures (Figure S2). In order to track QD-EphB2R movement relative to synapse position we expressed PSD95-GFP in neurons, to label the postsynaptic compartment, and imaged QD-EphB2Rs. Figure 3A shows a representative trajectory of a QD-EphB2R in relation to PSD-95. In this example, while the QD-EphB2R clearly diffuses into the synaptic compartment (blue), directed motion (red) is only seen outside synapses. Quantification of trajectory segments showing diffusion versus directed motion demonstrates that EphB2Rs with a velocity are very rarely seen at the synapse, compared with diffusing receptors (Figure 3B). These data indicate that surface EphB2Rs exhibit fast, directed motion along dendrites and between postsynaptic specializations.

**Figure 3.**
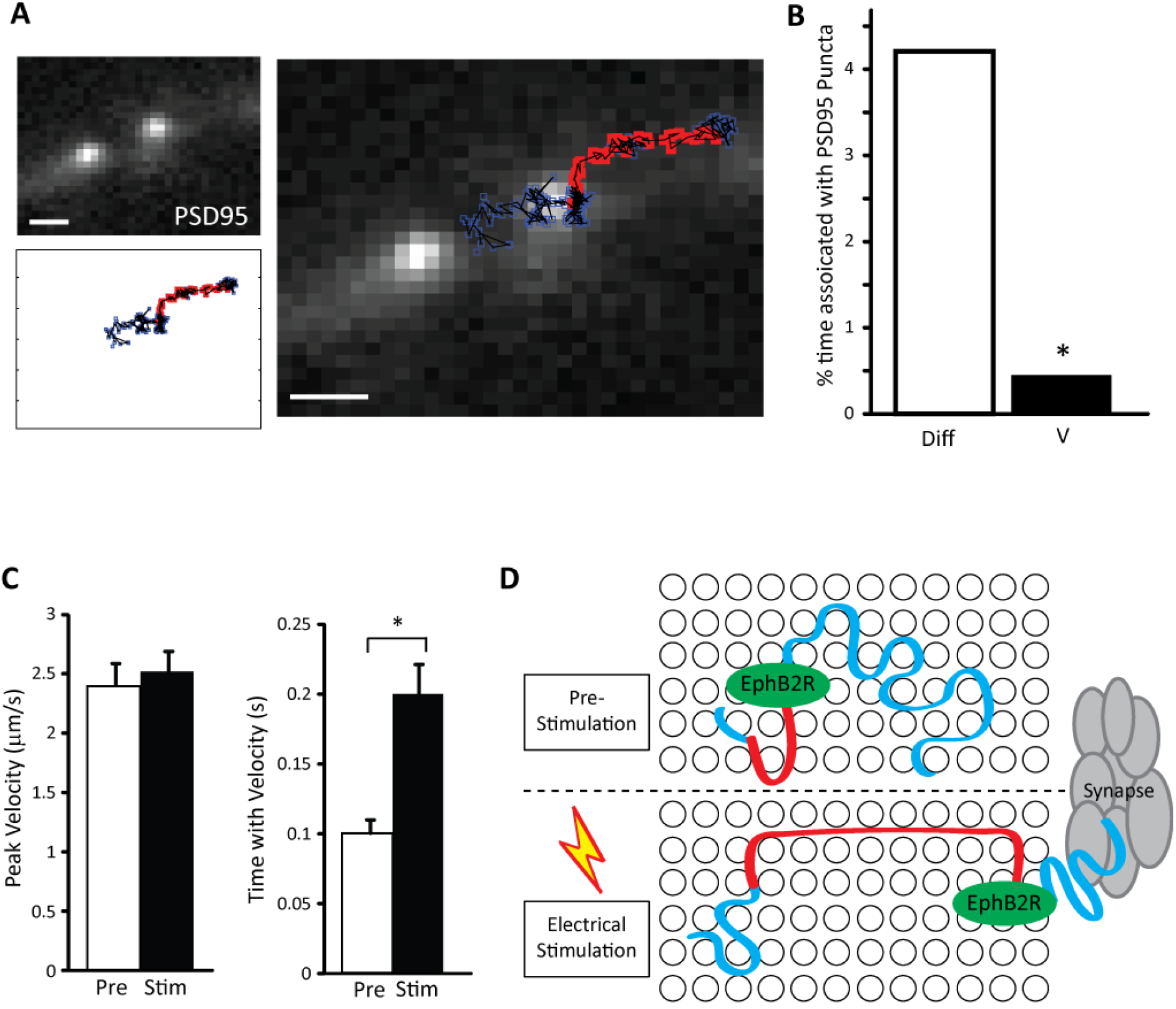
Directed motion is spatially restricted and activity dependent. (**A**) Representative trajectory of a single QD-EphB2R along a neuronal process expressing PSD-95-GFP to localize synapses (see also Figure S2). Regions of the trajectory in which the QD-EphB2R has an associated velocity/ displays directed movement are illustrated in red, with diffusive movement in blue, scale bar 1μm. (**B**) Percentage of time in which diffusive QD-EphB2Rs (Diff) or those displaying directed motion (V) are associated with PSD-95-GFP puncta (n=41, p<0.05, Chi-squared). (**C**) Field stimulation of the neuronal network increases time spent with a velocity, without changing the peak velocity. (**D**) Proposed model for the movement of QD-EphB2Rs in the membrane before and after electrical stimulation. QD-EphB2Rs (shown in green) diffuse in the cell membrane (blue tracks) displaying directed motion (red tracks) for short periods of time. QD-EphB2Rs located near to or at the synapse only display diffusive motion. After electrical stimulation, the directed motion of QD-EphB2Rs occurs across longer timescales.

Finally, in order to determine whether this directed movement might be regulated by neuronal activity, we analysed the directed motion characteristics of QD-EphB2Rs following electrical field stimulation. We found that QD-EphB2Rs travel for a longer duration in directed motion mode following stimulation with 900 action potentials at 20 Hz (mean duration pre stimulation: 0.09±0.05s, mean duration 5 minutes post stimulation: 0.14±0.09) without an associated change in peak velocity (peak velocity pre: 2.40±0.90μm/s, mean duration 5 minutes post stimulation: 2.52±0.73 μm/s) (Figure 3C). Thus, increased neuronal activity promotes rapid inter-synaptic movement of surface EphB2Rs.

## Discussion

We propose that cholesterol-dependent ‘conduits’ within the dendritic plasma membrane may allow brief spells of rapid surface receptor travel between synapses. Under conditions of increased neuronal activity receptors spend longer in these conduits (model illustrated in Figure 3D).

The dominant model of cell membrane structure remains the fluid mosaic model (Goni, 2014; Singer & Nicolson, 1972) where both proteins and lipids move within the fluid lipid bilayer, which is often described as having the consistency of olive oil (Edidin, 2003). This fluidity includes both translational and rotational (about their own axis) movement of molecules. In biological membranes, cholesterol is thought to ‘buffer’ fluidity, increasing fluidity at lower temperatures and reducing it at higher ones (Alberts, 2015; Maxfield & van Meer, 2010). Cholesterol is also associated with a liquid ordered phase where rotational movement is restricted (Goni, 2014), possibly in the context of lipid nanodomains, or ‘rafts’ (Simons & Ikonen, 1997; Simons & van Meer, 1988). Indeed, high resolution live cell imaging studies have indicated that cholesterol plays a role in ‘trapping’ sphingolipids and GPI-anchored proteins (Dietrich, Yang, Fujiwara, Kusumi, & Jacobson, 2002; Suzuki et al., 2012; Wenger et al., 2007). Consistent with this, our finding of significant increases in mean values of α for both super-confined and confined diffusing EphBRs, together with the overall increase in diffusion coefficient, suggest release from such domains. Although manipulation of cholesterol levels can also have effects on the actin cytoskeleton (Goodwin, Drake, Remmert, & Kenworthy, 2005; Kwik et al., 2003; Sun et al., 2007) including the organization of GPI-anchored proteins in the membrane (Goswami et al., 2008), and indeed actin-dependent, directional membrane flows have recently been described (Ashdown et al., 2017), the lack of effect on directed motion from modulation of the actin network argues that this is not actin-mediated, but rather dependent on the membrane itself. A possibility is that highly transient fluid currents may provide the conduits for directional movement of proteins in the neuronal membrane. In a polarized cell such as a neuron, with the need to rapidly move proteins in response to stimuli such as activity, this could represent an alternative and energy efficient method of protein delivery.

## Materials and Methods

### Hippocampal Cultures and Transfection

Dissociated hippocampal neuronal cultures were prepared from embryonic day 18 Sprague Dawley rats of both sexes in accordance with all institutional and national guidelines. Hippocampi were dissociated using trypsin (5 mg/ml for 15 min at 37°C; Worthington), triturated through narrow diameter Pasteur pipettes and plated at 350 cells/mm^2^ on glass coverslips pre-coated with poly-d-lysine (50 μg/ml; Sigma) and laminin (20 μg/ml). Neurons were incubated (37°C, 6% CO_2_) in a 50:50 mixture of neurobasal media supplemented with B27 (2%) and neurobasal media with fetal calf serum (2%), with additional glutamax (500 μM) and penicillin-streptomycin (100 μg/ml). At 7 days *in vitro* (DIV) half of this medium was replaced with neurobasal medium plus B27 supplement. Unless otherwise specified, all culture reagents were from Gibco.

Neurons (9-10 DIV) were transfected with PSD-95-GFP using Effectene Transfection Reagent (Qiagen) according to the manufacturer’s instructions.

### Live Single Molecule Optical Microscopy

Dissociated hippocampal neurons (10-14 DIV) were incubated for 10 minutes at 37° C in conditioned cell culture medium containing 1% casein and 1% pre-coupled QD-EphB2R mix. Pre-coupling was carried out by incubating QD 655 (Fisher) with EphB2R antibody (BD Pharmingen) in a 1:4 ratio in sterile PBS. Neurons were subsequently washed in Tyrode’s solution. Images were acquired <20minutes after the staining protocol was complete.

Images were acquired using an Olympus IX71 inverted microscope with a 100X, 1.4 NA oil immersion objective, coupled to a Photometrics Evolve EMCCD camera and associated Slidebook software (Intelligent Imaging Innovations, Denver). Illumination was provided by an LED emitting at 470nm. The filter cube used for QD imaging contained a dichroic mirror at 475nm and a 40nm emission filter centred at 655nm. Filters were purchased from Chroma Technologies. QD imaging movies were acquired with an acquisition rate of 32.9Hz and were 3000 frames in length. Tyrode’s solution contained: 128mM NaCl, 5mM KCl, 1mM MgCl_2_, 2mM CaCl_2_, 15mM HEPES, 4mM NaHCO_3_. Temperature was maintained at 35°C for the duration of the experiments with the aid of an objective warmer and heated chamber (Harvard Apparatus).

Electrical stimulation experiments were conducted using a field stimulation chamber (Harvard Apparatus) coupled to a SD9 stimulator delivering 900 pulses at 20Hz. Changes in the motion of QD-EphB2Rs were analysed 5 minutes after delivery of electrical stimulation.

### Filipin Fluorescence to measure Cholesterol Depletion

After treatment with methyl-β-cyclo-dextrin (1mM, 30mins at 37°C), dissociated hippocampal neurons (10-14DIV) were fixed in 4% paraformaldehyde for 10 minutes and permeabilised with 0.01% saponin . Cells were then incubated with filipin (Sigma) complex (100μg/ml) for five minutes and imaged immediately to avoid photo-bleaching.

Filipin fluorescence was measured using the ImageJ plugin: NeuronJ to trace neuronal processes, and then a custom written MATLAB (Mathworks) routine to measure the fluorescence intensity of the traced processes and subtract background from the images.

### Drugs and Reagents

Unless otherwise specified, all drugs were added to the recording chamber for 10 minutes prior to recording, dissolved in DMSO (final concentration 0.05%) and purchased from Sigma-Aldrich: methyl-β-cyclo-dextrin, filipin complex, jasplakinolide (Tocris), latrunculin A (Tocris), nocodazole and QSY-21 (Life Technologies). For experiments with low dose nocodazole (200nM) cells were incubated in the drug for 4 hours are 37°C prior to recording.

Pre-clustered ephrinB1-Fc was obtained by incubating ephrinB1-Fc (R&D Systems) with antibodies against Fc fragments (Sigma) in a 1:10 ratio for 1hour at 37°C. The mixture was then maintained until use at 0°C. Pre-clustered ephrinB1-Fc was applied to the cells at a final concentration of 2.2μg/ml. In control experiments Fc fragments replaced ephrinB1-Fc at an equivalent concentration.

### Receptor Tracking

The imaged molecules were detected and tracked in MATLAB using the plugin software UTrack 2.1 (Jaqaman et al., 2011). The individual QD-EphB2R tracks were then further analysed in MATLAB using custom written algorithms previously described (Arcizet et al., 2008). Briefly, the displacement between every pair of sub-resolution co-ordinates was analysed enabling the mean squared displacement (MSD) to be calculated and the trajectory motion mode to be assigned. We calculate time resolved MSD by calculating the local MSD function over 40 frames. The local MSD function is calculated according to:

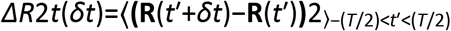

The diffusion coefficient (D) was calculated by a linear least squares regression fit of the first three points of this function.

The motion mode is determined by fitting the power law and calculating α over the first twenty points of the MSD function:

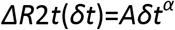

QD-EphB2Rs were determined to have a velocity when two criteria were satisfied: α greater than the critical value (1.561) and the statistical assessment of a polynomial fit to the MSD function was significantly better than a linear fit. The critical value for α was determined as being significantly greater than the α calculated for simulated random motion and is in agreement with Caspi and colleagues where a value of α>1.5 is synonymous with super diffusion (Caspi, Granek, & Elbaum, 2000). Instantaneous velocity (V) was calculated for segments of a trajectory where α> 1.561 for at least two time steps.

The time resolved MSD was calculated using sliding windows of 40 steps to enable extrapolation of instantaneous motion characteristics. This enabled automatic trajectory analysis of motion modes without *a priori* segmentation.

To compare experimental data with simulated, we used trajectories without blinks and simulated 350 trajectories of similar lengths displaying random motion. The simulated trajectories were then analysed using time resolved MSD analysis and the distribution of instantaneous α obtained. The values of instantaneous α obtained for random simulated trajectories was 0.928±0.248.

### Data and Statistics

Unless stated otherwise in the text, all statistics are presented as the mean ± standard deviation. Statistical analysis was carried out using the Student t test, ANOVA or Chi-squared test where appropriate.

## Acknowledgments

The help of Daniel Choquet and the Bordeaux Imaging Center, part of the national infrastructure France BioImaging, granted by ANR-10INBS-04-0, is acknowledged. This research was supported by BBSRC grant BB/P000479/1 (to L.C.A.), MRC grant G0802613 (to N.J.E.) and the Sackler Institute for Translational Neurodevelopment (R.D.T.). L.C.A. is a NARSAD Young Investigator. We thank Uwe Drescher (KCL, London) for the kind gift of Fc fragments, ephrinB1-Fc and antibodies against Fc, Oscar Marin and members of the Andreae and Emptage labs for critical discussions.

## Movie

### Quantum Dot labelled EphB2Rs moving on hippocampal neurons

Time-lapse imaging of QD-EphB2Rs (red dots) superimposed on brightfield image of dissociated hippocampal neurons. White arrow indicates stationary QD-EphB2R, red arrow indicates QD-EphB2R that displays different patterns of movement during course of imaging.

